# Timing matters: Sex differences in acute and chronic outcomes following repetitive blast mild traumatic brain injury

**DOI:** 10.1101/2022.10.11.511013

**Authors:** Britahny M. Baskin, Aric F. Logsdon, Suhjung Janet Lee, Brian D. Foresi, Elaine Peskind, William A. Banks, David G. Cook, Abigail G. Schindler

## Abstract

**Background:** Repetitive blast-related mild traumatic brain injury (mTBI) caused by exposure to high explosives is increasingly common among warfighters as well as civilians. While women have been serving in military positions with increased risk of blast exposure since 2016, there are few published reports examining sex as a biological variable in models of blast mTBI, greatly limiting diagnosis and treatment capabilities. As such, here we examined acute and chronic outcomes of repetitive blast trauma in female and male mice in relation to potential behavioral, inflammatory, microbiome, and vascular dysfunction.

**Methods:** In this study we utilized a well-established blast overpressure model to induce repetitive (3x) blast-mTBI in both female and male mice. Acutely following repetitive exposure, we measured serum and brain cytokine levels, blood-brain barrier (BBB) disruption, fecal microbial abundance, and locomotion and anxiety-like behavior in the open field assay. Chronically, in female and male mice we assessed behavioral correlates of mTBI and PTSD-related symptoms commonly reported by Veterans with a history of blast-mTBI using the elevated zero maze, acoustic startle, and conditioned odorant aversion paradigms.

**Results:** Repetitive blast exposure resulted in both similar and disparate patterns of acute serum and brain cytokine as well as gut microbiome changes in female and male mice. Acute BBB disruption following repetitive blast exposure was apparent in both sexes. While female and male blast mice both exhibited acute locomotor and anxiety-like deficits in the open field assay, only male mice exhibited chronic adverse behavioral outcomes.

**Discussion:** Representing a novel survey of potential sex differences following repetitive blast trauma, our results demonstrate unique similar yet divergent patterns of blast-induced dysfunction in female vs. male mice and highlight novel targets for future diagnosis and therapeutic development.

## INTRODUCTION

Traumatic brain injury (TBI) is currently a leading cause of death and disability not just in the United States but globally (Johnson & Griswold, 2017; Maas et al., 2017; Taylor, 2017). Affecting every segment of the population, TBI is often associated with significantly decreased quality of life and increased financial burden (Di Battista et al., 2012; Malec et al., 2017; Ozga et al., 2018; Taylor, 2017). Post-TBI symptomatology can be highly variable and comorbid with other diagnoses such as posttraumatic stress disorder (PTSD) and chronic pain, resulting in limited diagnosis and treatment capabilities. The vast majority of preclinical TBI research has focused only on male research animals, which may express different symptom trajectories than female animals, resulting in a critical knowledge gap in the field at a time when women are at increasingly high risk for repetitive TBI exposure (Giordano et al., 2020; Gupte et al., 2019; McCabe & Tucker, 2020; Späni et al., 2018).

Blast overpressure (BOP) waves, such as those caused by improvised explosive devices (IEDs) and industrial accidents, are becoming an increasingly common cause of TBI. Referred to as the “signature injury” of the conflicts in Iraq/Afghanistan (OEF/OIF/OND), repetitive blast exposure is the primary source of mTBI in warfighters, a significant driver of comorbid PTSD, and a major source of morbidity among Veterans enrolled in the VA health care system (Adamson et al., 2008; Hendrickson et al., 2018; O’Neil et al., 2013; Owens et al., 2008; Wenger et al., 2018). In these conflicts, an estimated 75% of all mTBI reported by Servicemembers are a result of blast exposure caused by detonation of high explosives (Adamson et al., 2008; Owens et al., 2008). Furthermore, multiple deployments are common (2.77 million Servicemembers have served on 5.4 million deployments since 2011), resulting in a high potential for repetitive blast exposures (Agoston, 2017; Carr et al., 2016; Ravula et al., 2022; Simmons et al., 2020). Currently, there is a lack of research on how these injuries may differentially impact people identifying as male vs. female, yet females currently represent ∼15% of active duty Servicemembers and ∼20% of the United States Reserve and Guard (Dye et al., 2016; Iverson et al., 2011; Kamarck, 2015; Street et al., 2013). Indeed, few preclinical studies have included both male and female animals and the few that have, have exclusively focused on impact TBI, a single TBI exposure, and/or only examined acute timepoints. While a handful of reports have characterized potential sex differences following a single blast exposure (Hubbard et al., 2022; Kawa et al., 2020; McNamara et al., 2022; Russell, Handa, et al., 2018; Russell, Richardson, et al., 2018), very little is known about the effects of repetitive blast exposure in male vs. female rodents.

Given the lack of preclinical research using models of repetitive blast exposure in female rodents, this study was envisioned as a novel survey of adverse outcomes commonly seen following repetitive blast trauma in warfighters, with a specific focus on delineating acute and chronic outcomes related to inflammatory, blood-brain barrier (BBB), microbiome, and behavioral pathology. Results highlight both similar and disparate outcomes in female vs. male mice at acute timepoints and an effect only in male mice at chronic time points following repetitive blast mTBI. Together, these results highlight new targets for diagnosis and treatment development and confirm the need for increased research dedicated to understanding how repetitive blast trauma affects diverse populations.

## MATERIALS AND METHODS

### Animals

All experiments utilized female and male (as determined by genital appearance at weaning) C57Bl/6 mice aged 9-11 weeks old at time of arrival to VA Puget Sound. Mice were housed three per cage by sex on a 12:12 light:dark cycle (lights on at 06:00), and were given *ad libitum* food and water. All animal experiments were carried out in accordance with Association for Assessment and Accreditation of Laboratory Animal Care guidelines and were approved by the VA Puget Sound Institutional Animal Care and Use Committees. Mice were acclimated to the housing room for a week following arrival and subsequently handled for an additional week prior to sham or blast exposure. Three experimental timelines were employed (Figure 1a) in separate sets of mice. To increase rigor and reproducibility, each experimental timeline included at least two cohorts of mice each run at separate times.

**Figure 1.**
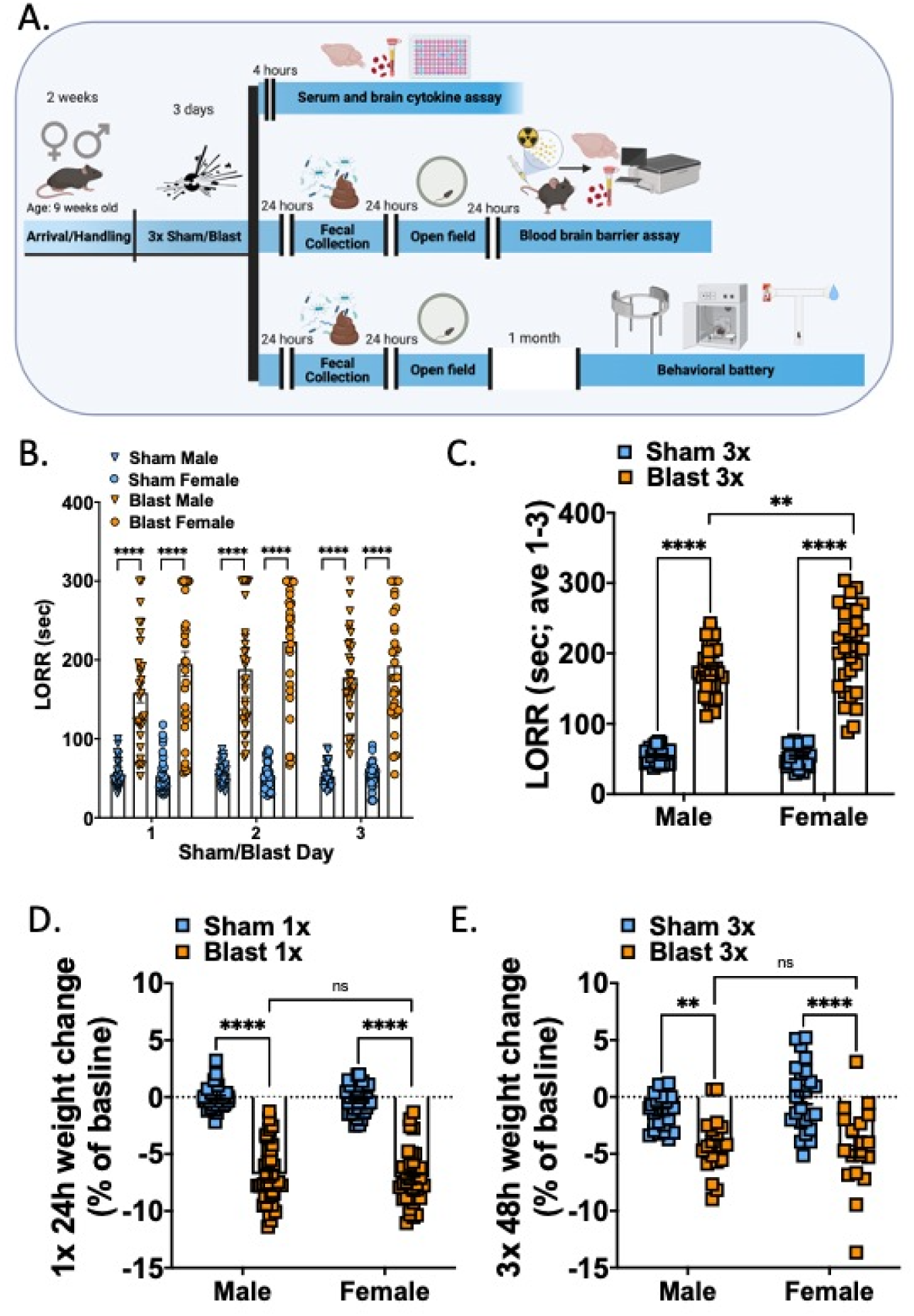
Female and male mice both display increased LORR and weight loss acutely following repetitive blast exposure. (A) Experimental timelines. Three groups of female and male mice were used. (B) LORR is increased acutely following each blast exposure in female and male mice. (C) Mean LORR across days 1-3 of blast exposure is increased in female and male mice and female blast mice are significantly higher than male blast mice. (D) Both female and male mice lose weight to a similar extent 24 hours following a single blast exposure. (E). Both female and male mice lose weight to a similar extent 48 hours following final repetitive (3x) blast exposure. Two-way ANOVA, Šídák’s multiple comparison post-hoc. **p ≤ 0.01, ****p ≤ 0.0001, ns = not significant. Values represent mean ± SEM.

### Model of Blast Trauma

The shock tube (Baker Engineering and Risk Consultants, San Antonio, TX) was designed to generate blast overpressures that mimic open field high explosive detonations encountered by military Servicemembers in combat. Design and modeling characteristics have been described in detail elsewhere (Baskin et al., 2021; Huber et al., 2013; Logsdon et al., 2020; Meabon et al., 2016; Schindler et al., 2017; Schindler, Terry, et al., 2021; Schindler, Baskin, et al., 2021). Briefly, mice were weighed and then anesthetized with isoflurane (induced at 5% and maintained at 2-3% for the duration of the blast), secured against a gurney, and placed into the shock tube oriented perpendicular to the oncoming blast wave (ventral body surface towards the blast). Utilizing condensed helium, pressurized air is built up on one end of the tube and released in a way that creates a blast overpressure wave that induces neuropathological and behavioral changes in line with blast mild traumatic brain injuries (mTBI) in warfighters (Ghai et al., 2020; Meabon et al., 2020; Peskind et al., 2011; Petrie et al., 2014; Schindler, Baskin, et al., 2021; Schindler et al., 2017; Wilkinson et al., 2012). Sham (control) animals received anesthesia only for a duration matched to blast animals. Repeated blast/sham exposures occurred successively over the course of three days (one per day). The blast overpressure (BOP) peak intensity (psi), initial pulse duration (ms), and impulse (psi▪ms) used were in keeping with mild blast injury (19.1 +/- 0.09 psi). Under these experimental conditions, the overall survival rate exceeded 97%, with blast-exposed mice presenting comparable to sham-exposed mice by inspection 2-4 hours-post blast exposure as previously reported (Baskin et al., 2021; Huber et al., 2013, 2016; Logsdon, Meabon, et al., 2018; Logsdon et al., 2020; Meabon et al., 2016; Schindler et al., 2017; Schindler, Terry, et al., 2021; Schindler, Baskin, et al., 2021). Following sham/blast exposure and removal from isoflurane, loss of righting reflex (LORR) was recorded as the amount of time it took for animals to right themselves. Once animals were able to regain full sternal recumbency, they were weighed and then returned to their cages for recovery.

### Cytokine Measurement

Mice were euthanized via cervical decapitation 4 hours after their last sham or blast exposure. Trunk blood was collected in 1.5-mL capacity serum-separator tubes, allowed to clot at room temperature for 30-40 minutes, and then centrifuged at 3,000 x g for 10 minutes. Serum was then aliquoted and stored at -80°C until analyzed. Whole brains were also collected at the time of euthanasia, hemisectioned, and flash frozen at -80°C. One hemisphere of brain tissue was then lysed in a 0.02% Triton-X homogenization buffer of 10mM HEPES, 1.5mM MgCl2, and 10mM KCl, with fresh protease/phosphatase inhibitor cocktail (Sigma) in a bead homogenizer. The samples were then centrifuged at 4°C at 18,000 x g for 10 minutes and the supernatant was aliquoted and stored at -80°C. Pro- and anti-inflammatory cytokine level were then analyzed using the IDEXX (Columbia, MO) Cytokine Mouse 25-Plex Panel and values were normalized by protein concentration quantified with a BCA Protein Assay Kit.

### Fecal Microbiome

Fresh fecal pellets were collected 24 hours following final sham or blast exposure. Collection occurred in the morning between 09:00 and 11:00 using sterile technique. Fecal pellets were flash frozen in liquid nitrogen and stored at -80°C until shipment to Diversigen for downstream processing and analysis. Briefly, the DNA of fecal pellets were extracted and sequenced by Diversigen Inc. (New Brighton, MN) using their BoosterShot Shallow Shotgun Sequencing. DNA sequences were aligned to a curated database containing all representative genomes in RefSeq for bacteria. Alignments were made at 97% identity against all reference genomes. Every input sequence was compared to every reference sequence in the Diversigen Venti database using fully gapped alignment with BURST. For taxonomy assignment, each input sequence was assigned the lowest common ancestor that was consistent across at least 80% of all reference sequences tied for best hit. Samples with fewer than 10,000 sequences were discarded. Operational taxonomic units (OTUs) accounting for less than one millionth of all strain-level markers and those with less than 0.01% of their unique genome regions covered (and < 0.1% of the whole genome) at the species level were discarded. The number of counts for each OTU was normalized to the OTU’s genome length.

### Blood-brain barrier disruption assessment

Following established procedures (Banks et al., 2015; Logsdon, Meabon, et al., 2018; Logsdon et al., 2020), albumin (Sigma, St. Louis MO), (∼66.44 kDa), a blood-borne molecule with minimal penetration into the CNS, was labeled with 99mTc (GE Healthcare, Piscataway, NJ). A mixture of 240 mg/ml stannous tartrate and 1 mg/ml albumin was adjusted to pH 3.0 with HCl and one millicurie of 99mTc-NaOH4 was added to this mixture and incubated for 20 min. The ^99m^Tc-albumin was purified on a column of G-10 Sephadex (GE Healthcare) in 0.1 ml fractions of phosphate buffer (0.25 M). Radioactivity in the purified ^99m^Tc-albumin peak was more than 90% acid precipitable in an equal volume of 1% bovine plasma albumin (BSA) and trichloroacetic acid (30%). 5 × 10^6^ cpm/mouse of purified ^99m^Tc-albumin fraction was prepared in a final volume (0.2 ml/mouse) of lactated Ringer’s solution containing 1% BSA.

In keeping with previous experiments (Logsdon, Meabon, et al., 2018; Logsdon et al., 2020), 72 hours after the final blast/sham exposure, mice were anesthetized with urethane (4 g/kg; 0.2 ml; ip), the jugular veins were exposed, and ^99m^Tc-albumin (5 × 10^6^ Counts per minute (cpm)) in 0.2 ml of lactated Ringer’s solution with 1% BSA was injected into the jugular vein and allowed to circulate for ten minutes. After ten minutes of circulation, the abdominal aorta was clamped with hemostats, severed, and blood collected in 1.5-mL capacity tubes containing heparin and then centrifuged at 4 degrees C at 3,200 x g for 10 minutes. Plasma was then aliquoted and stored at -80°C until analyzed. Immediately after blood collection, the vascular space of the brain was perfused with 20 ml of lactated Ringer’s solution through the left ventricle of the heart in less than 1 min, thus washing out the vascular contents of the brain so that only ^99m^Tc-albumin which had leaked into the brain remained. The brain was then collected and flash frozen in liquid nitrogen. ^99m^Tc radioactivity in the plasma and brain was measured by a gamma counter. Brain tissue radioactivity was calculated by dividing the cpm in the brain by the weight of the brain to yield cpm/g. Plasma radioactivity was calculated by dividing the cpm in the plasma by the microliters of plasma counted to yield cpm/microliter. The brain tissue radioactivity was then divided by the corresponding plasma radioactivity and the results given in units of microliters/gram of brain tissue.

### Behavioral assays

Acute (48 hours post): To probe potential locomotor deficits and anxiety-like behavior acutely following sham/blast exposure, mice were tested in an open field assay. Mice were allowed 5 minutes to explore a large circular open space (1 meter diameter) and their movements were recorded from above and analyzed with Anymaze (Wood Dale, IL). On this test, decreases in the amount of time spent in the middle of the environment is indicative of an anxiety-like phenotype. The total distance traveled, the delay to first enter the center of the field, time spent in the center, and entries into the center were recorded and analyzed.

Chronic (1 month post): A behavioral battery consisting of three testing paradigms was conducted over 1 week (one test paradigm per day) starting one month after the last sham/blast exposure. The order of behavioral tests was specifically chosen to go from the least stressful to the most stressful task to prevent carryover distress from one behavior to the next.

*Elevated zero maze (EZM)*: Animals were allowed to explore an elevated zero maze (Maze Engineers, Skokie, IL) for 5 min. Decreased time spent exploring the open arms is thought to reflect an anxiety-like behavior. Movement was recorded from above and analyzed using Anymaze (Stoelting, Wood Dale, IL).

*Acoustic startle (AS):* In accordance with previous reports (Baskin et al., 2021) acoustic startle habituation and prepulse inhibition were measured using SR-LAB acoustic startle boxes (San Diego Instruments, San Diego, CA). Following a 5-min acclimation period, startle habituation testing consisted of 50 trials of 120-dB pulses delivered with an inter-trial interval of 7–23 s. Following a two min break period, prepulse inhibition (PPI) was next assessed and consisted of forty trials of 81-dB prepulse followed by a 120-dB pulse with varying interstimulus interval (ISI) of 2–1,000 ms (five trials each). Due to the established effects of blast exposure on hearing loss, we use these established within-subject analysis approaches of startle habituation and PPI in attempts to mitigate potential confounds of hearing loss on startle outcome measures and interpretation.

*Conditioned odorant aversion (COA)*: As previously described (Schindler et al., 2021), mice were first exposed to a neutral almond scent starting five minutes prior to sham/blast exposure by placing a mesh tea ball containing a quarter nestlet with 20 ul almond extract into the home cage. The tea ball and the nestlet with almond scent was refreshed daily and remained in the home cage until 24 hours after the final sham/blast exposure. To measure blast-induced aversion at the chronic testing time point, a tea ball with almond extract nestlet was placed in the left arm of a tea maze and a tea ball with a clean nestlet was placed in the right arm of the t-maze. Animals were then placed in the long arm of the T-maze, equidistant away from either tea ball and given 5 min to explore the entire maze. Latency to enter and time spent in each of the two distal ends of the short arms was recorded and analyzed using Anymaze (Stoelting, Wood Dale, IL).

### Analysis

Data are expressed as means ± SEM. Differences between groups were determined using two-way (between-subjects design: sex and exposure factors) analysis of variance (ANOVA) followed by posthoc testing using Sidak’s multiple comparisons test. Microbiome bioinformatics (alpha diversity (Shannon), beta diversity (Bray Curtis), and relative abundance) were performed by Diversigen. Shannon diversity and relative abundance at the taxonomic order level were compared across all groups using Kruskal-Wallis test followed by Mann-Whitney *U* test for individual group by group comparison if Kruskal-Wallis was significant at false discovery rate (FDR) p<0.1. Bray Curtis dissimilarity was used to estimate beta diversity among samples and represented using PCoA plot. Analysis of similarities (ANOSIM) was used to compare beta diversity across groups (999 permutations). Spearman correlation with FDR correction was used to examine associations between microbial abundance and behavioral parameters.

Hierarchical clustering was performed using Ward agglomeration method. Differentially expressed taxa were identified using linear discriminant analysis effect size (LEfSe) using the Huttenhower Galaxy module (Segata et al., 2011). Reported p values denote two-tailed probabilities of p ≤ 0.05 and non-significance (n.s.) indicates p > 0.05. Statistical analysis and visualization were conducted using Graph Pad Prism 4.0 (GraphPad Software, Inc., La Jolla, CA) and with custom Python scripts.

## RESULTS

### Repetitive blast exposure increases loss of righting reflex and acute weight loss in both female and male mice

Adult female and male mice were exposed to repetitive BOPs (1 BOP/day for 3 consecutive days) using a pneumatic shock tube as previously described (Huber et al., 2013; Logsdon, Meabon, et al., 2018; Schindler et al., 2017). Loss of righting reflex (LORR; the time it takes for a mouse to right itself following sedation) was measured starting immediately upon removal from isoflurane and consisted of placing the mouse on its back and waiting until it rights itself two times within 15 seconds. Following LORR, mice were weighed and then returned to their home cage to recover.

Repetitive blast exposure resulted in increased LORR in both male and female mice (Fig 1b) immediately following each sham/blast exposure (two-way RM ANOVA: main effect of exposure *F*(3,124) = 171.5, *p*<0.0001), but was not affected by sham/blast day (two-way RM ANOVA: main effect of time *F*(2,217) = 3.19, ns) and there was no interaction effect (two-way RM ANOVA: main effect of time *F*(6,248) = 1.30, ns). When examining the mean LORR across all three exposure days, there was a significant interaction effect between exposure and sex (two-way ANOVA: interaction effect *F*(1,124) = 6.674, *p*=0.01) and main effects of exposure (two-way ANOVA: main effect of exposure *F*(1,124) = 482.6, *p*<0.0001) and time (two-way ANOVA: main effect of time *F*(1,124) = 5.384, *p*=0.022). Šídák’s multiple comparison post-hoc test revealed that male blast mice (n=32) took longer to right themselves when compared to male sham mice (n=33; *p*<0.0001) and female blast mice (n=27) had longer LORR than both female shams (n=38; *p*<0.0001) and male blast mice (*p<*0.01).

Acute and sub-acute weight loss was also seen in both male and female blast mice (Fig 1d, e). Following one blast (Fig 1d), blast mice lost more weight (represented as a decrease in percent of their total baseline weight) than their sham controls (two-way RM ANOVA: main effect of exposure *F*(1,144) = 466.5, *p*<0.0001). There was no significant interaction effect between exposure and sex (two-way ANOVA: interaction effect *F*(1,144) = 0.07, ns) or main effect of sex (two-way ANOVA: main effect of sex *F*(1,144) = 0.346, ns). Šídák’s multiple comparison post-hoc found that both male and female blast mice lost weight when compared to their sham controls but there was no difference between male and female blast mice. When examining weight loss at 48 hours post (day of open field testing), there remained a main effect of blast (two-way ANOVA: main effect of exposure *F*(1,84) = 39.72, *p*<0.0001) and no significant interaction effect between exposure and sex (two-way ANOVA: interaction effect *F*(1,84) = 1.45, ns) or main effect of sex (two-way ANOVA: main effect of sex *F*(1,84) = 0.927, ns) (Fig 1e). Šídák’s multiple comparison post-hoc found that both male and female blast mice lost weight when compared to their sham controls but there was no difference in percent weight loss between male and female blast mice.

### Repetitive blast exposure differentially affects acute cytokine changes in serum and brain of female and male mice

To determine if acute changes in serum and brain cytokine levels were different between female and male mice following repetitive blast, we euthanized a subgroup of mice (Fig 1a) 4 hours following their final sham/blast exposure. Analysis revealed both blast and sex differences dependent on cytokine sample type (Fig 2-3). Results from two-way ANOVA (exposure, sex factors) followed by Šídák’s multiple comparisons tests for each analyze measured in appreciable concentration are reported in Tables 1 (serum) and 2 (brain).

**Figure 2.**
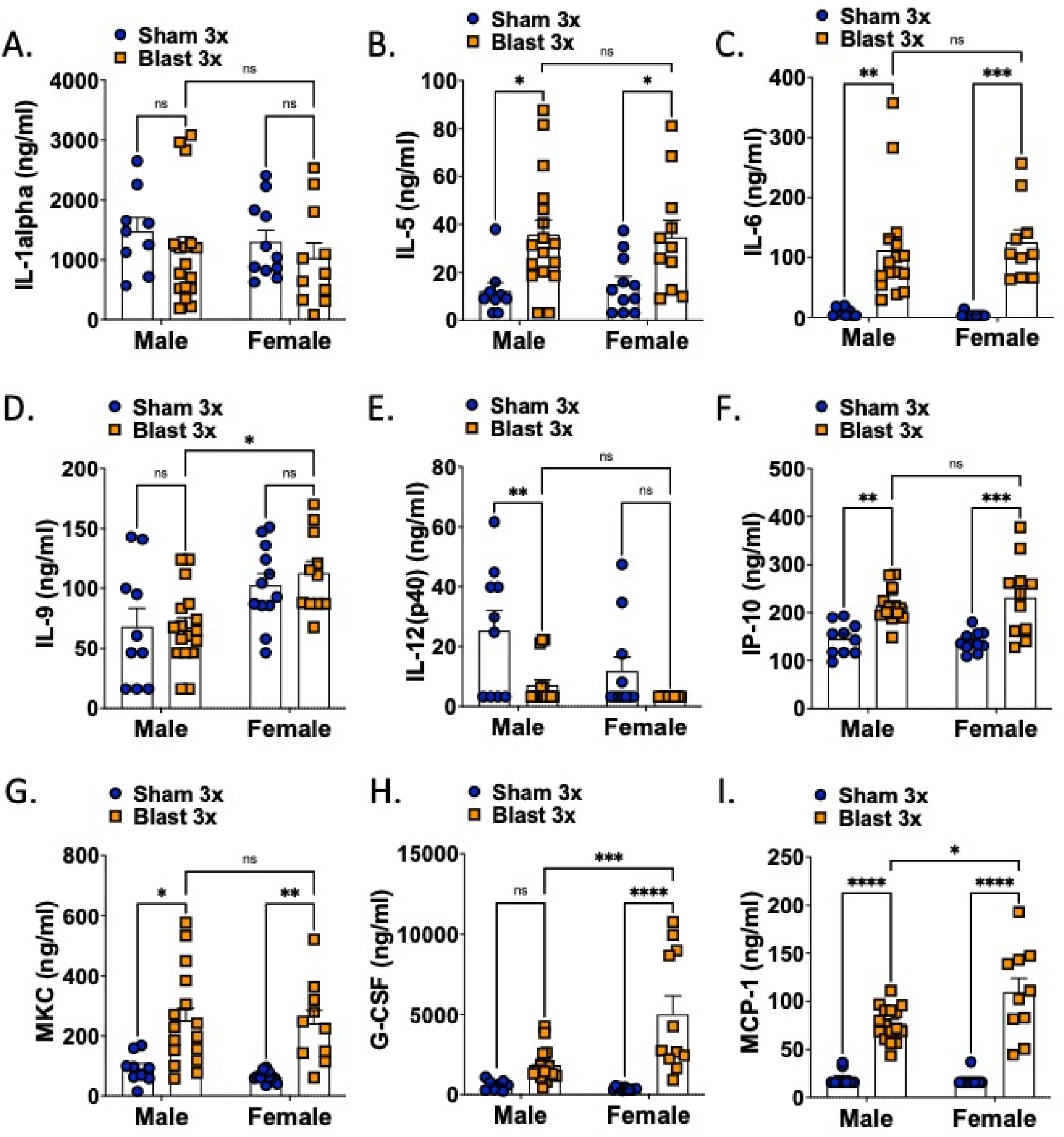
Serum cytokine levels are acutely affected by blast in both female and male mice. Two-way ANOVA Šídák’s multiple comparison post-hoc. *p ≤ 0.05, **p ≤ 0.01, ***p ≤ 0.001, ****p ≤ 0.0001, ns = not significant. Values represent mean ± SEM.

**Figure 3.**
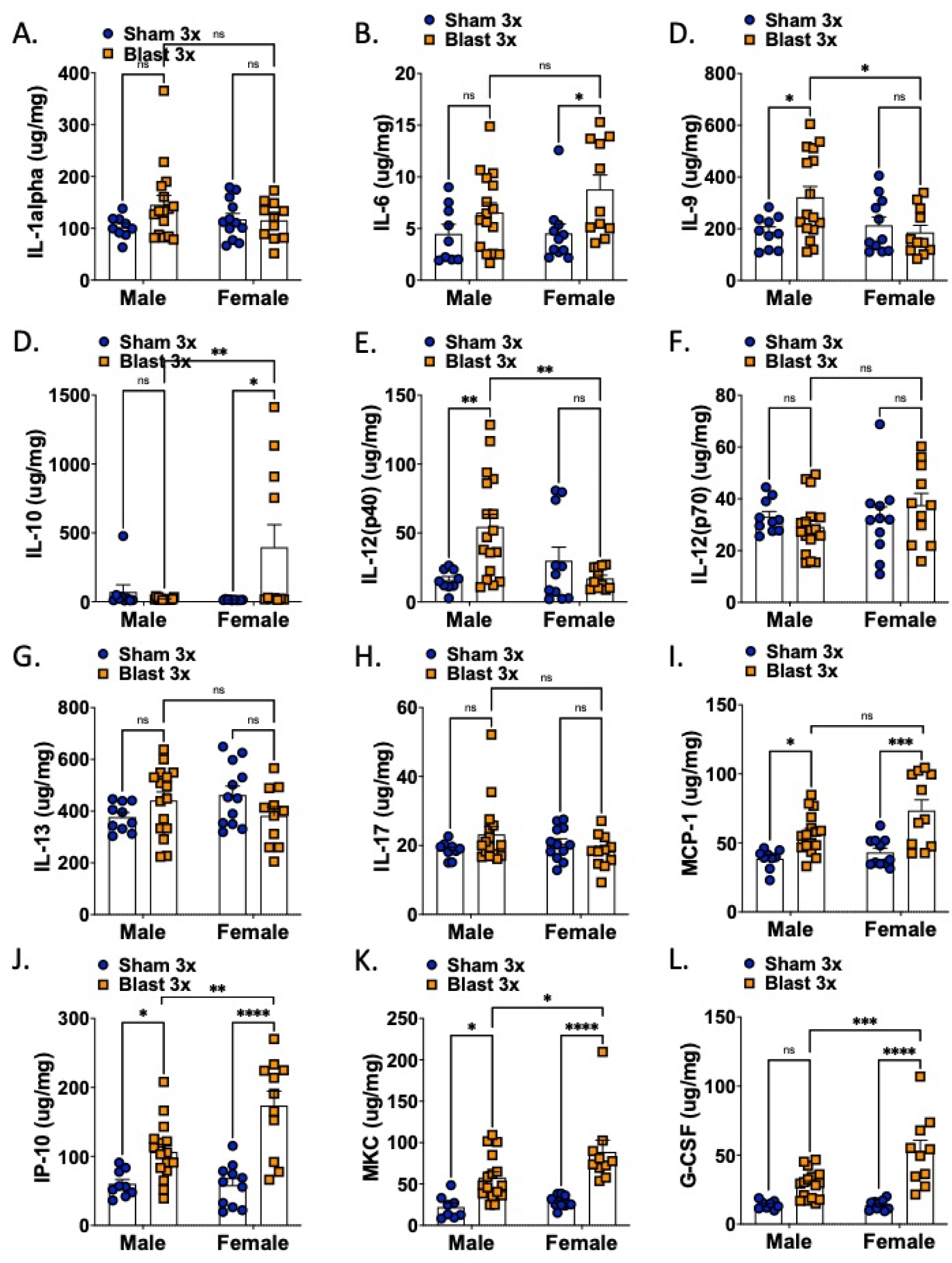
Brain cytokine levels are acutely affected by blast in both female and male mice. Two-way ANOVA Šídák’s multiple comparison post-hoc. *p ≤ 0.05, **p ≤ 0.01, ***p ≤ 0.001, ****p ≤ 0.0001, ns = not significant. Values represent mean ± SEM.

**Table 1.**
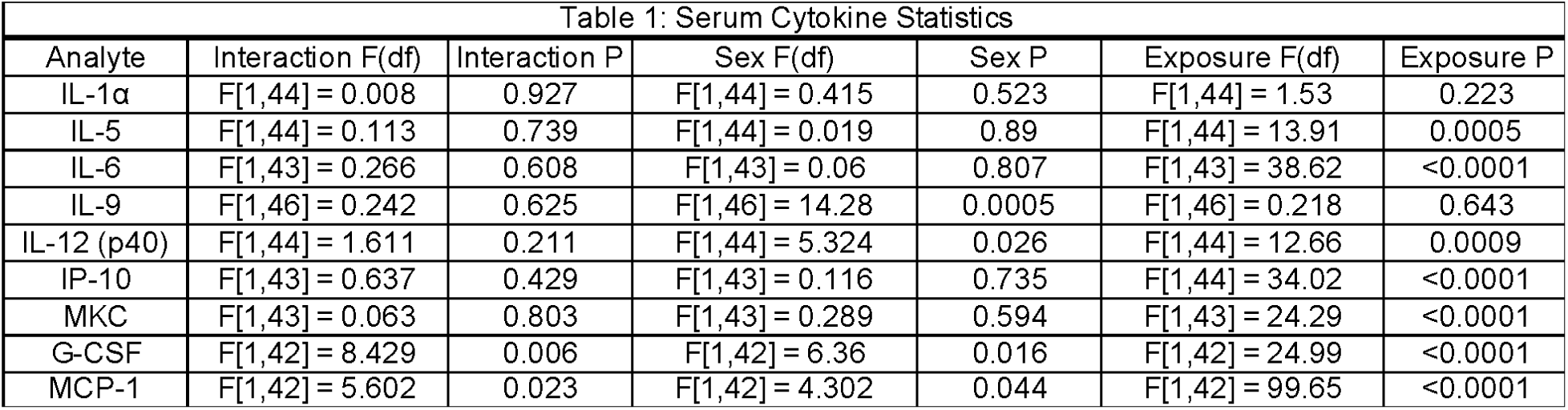
Statistical analysis of cytokine levels in blood serum four hours after the final blast. All analyses used two-way ANOVA and Šídák post-hoc analysis.

In the serum (Fig 2, Table 1), there was a main effect of blast on IL-6, G-CSF, IP-10, MKC, and MCP-1. There was main effect of sex on G-CSF and MCP-1. Only G-CSF and MCP-1 showed an interaction effect. Blast male and female mice only significantly differed in two cytokines, G-CSF and MCP-1, with blast males expressing significantly lower levels than blast females. In samples taken from whole brain (Fig 3, Table 2), there was a main effect of blast on IL-6, G-CSF, IL-10, MKC, and MCP-1. There were main effects of sex on G-CSF, IP-10, MKC, and MCP-1 and an interaction effect on IL-9, IL-10, IL-12(p40), IL-13, G-CSF, and IP-10. Blast males showed higher levels of IL-9 and IL-12(p40) as compared to blast females whereas blast females showed higher levels of IP-10, G-CSF, IP-10, and MKC as compared to blast males.

**Table 2.** Statistical analysis of cytokine levels in brain lysate four hours after the final blast. All analyses used two-way ANOVA and Šídák post-hoc analysis.

| Table 2: Brain Cytokine Statistics |  |  |  |  |  |  |
| --- | --- | --- | --- | --- | --- | --- |
| Analyte | Interaction F(df) | Interaction P | Sex F(df) | Sex P | Exposure F(df) | Exposure P |
| IL-1α | F[1,44] = 2.212 | 0.144 | F[1,44] = 0.249 | 0.62 | F[1,44] = 1.850 | 0.1807 |
| IL-6 | F[1,43] = 1.040 | 0.314 | F[1,43] = 1.107 | 0.299 | F[1,43] = 8.612 | 0.005 |
| IL-9 | F[1,44] = 5.247 | 0.027 | F[1,44] = 2.422 | 0.127 | F[1,44] = 2.256 | 0.14 |
| IL-10 | F[1,42] = 6.909 | 0.012 | F[1,42] = 3.38 | 0.062 | F[1,42] = 3.999 | 0.052 |
| IL-12 (p40) | F[1,44] = 9.873 | 0.003 | F[1,44] = 2.02 | 0.162 | F[1,44] = 2.48 | 0.123 |
| IL-12 (p70) | F[1,45] = 1.706 | 0.198 | F[1,45] = 1.143 | 0.291 | F[1,45] = 0.039 | 0.844 |
| IL-13 | F[1,45] = 4.977 | 0.031 | F[1,45] = 0.157 | 0.694 | F[1,45] = 0.061 | 0.805 |
| IL-17 | F[1,44] = 3.216 | 0.08 | F[1,44] = 0.619 | 0.436 | F[1,44] = 0.363 | 0.55 |
| MCP-1 | F[1,43] = 1.585 | 0.215 | F[1,43] = 4.919 | 0.032 | F[1,43] = 25.01 | <0.0001 |
| IP-10 | F[1,43] = 6.567 | 0.014 | F[1,43] = 5.877 | 0.02 | F[1,43] = 36.15 | <0.0001 |
| MKC | F[1,42] = 2.2 | 0.146 | F[1,42] = 5.502 | 0.031 | F[1,42] = 33.37 | <0.0001 |
| G-CSF | F[1,41] = 8.167 | 0.007 | F[1,41] = 7.932 | 0.007 | F[1,41] = 40.83 | <0.0001 |

### Unique gut microbiome signatures in female vs. male mice following repetitive blast exposure

To determine if repetitive blast exposure differentially affected the gut microbiome in female vs. male mice, fecal samples were collected 24 hours post final sham/blast exposure and analyzed using shallow shotgun sequencing. Alpha diversity measured using the Shannon Index (Figure 4a) was not different when comparing across groups (Kruskal Wallis *F*(4,94) = 2.47, *p*=0.57). Conversely, beta diversity measured using Bray Curtis distance (Figure 4b) was significantly different across groups (ANOSIM, 999 permutations, *p*=0.001). Furthermore, we found significant differences in abundance of nine bacteria orders (Figure 4c). Finally, we examined relative abundance differences at the species level using LeFSe (Huttenhower Lab Galaxy) and found unique species characterizing each of the four experimental groups (Figure 5).

**Figure 4.**
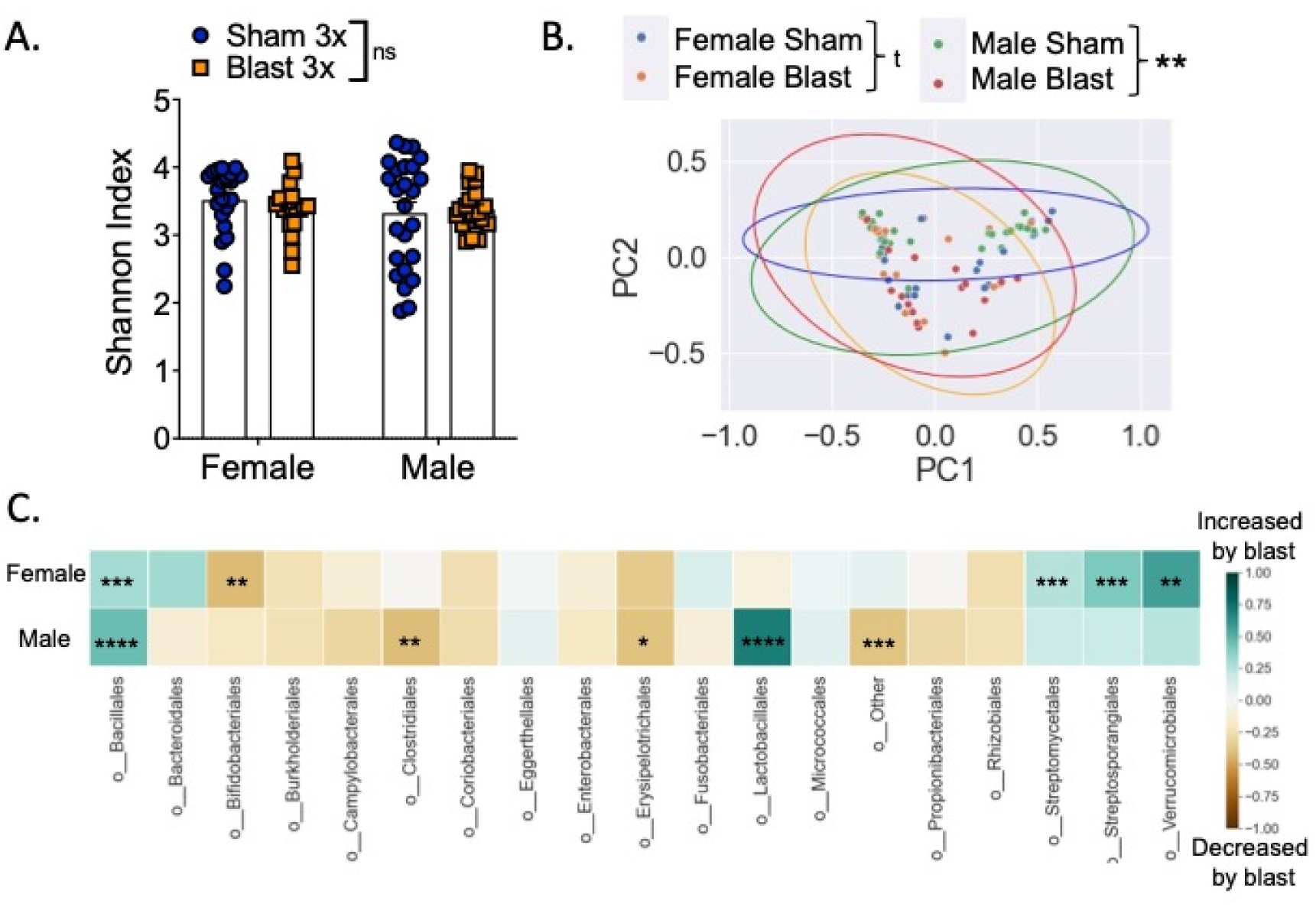
Differential effects of repetitive blast on acute gut microbiome changes in female vs. male mice. (A). No difference in Shannon (alpha) diversity as a function of exposure type or sex. (B) PCoA on Bray-Curtis dissimilarity distances (beta diversity) among the four groups examined. Each point represents an individual sample colored according to group. Ellipses represent 95% CI around cluster centroid for each experimental group. (C) Differences between female sham vs. blast mice (top row) and male sham vs. blast mice (bottom row) at the order level, expressed as mean relative abundance z score (z score computed separately for female and male mice). Kruskal-Wallis test followed by Mann-Whitney *U* test for individual group by group comparison if Kruskal-Wallis was significant at FDR p<0.1 (A, C). Analysis of similarities (ANOSIM) (B). *p ≤ 0.05, **p ≤ 0.01, ***p ≤ 0.001, ****p ≤ 0.0001, ns = not significant. Values represent mean ± SEM.

**Figure 5.**
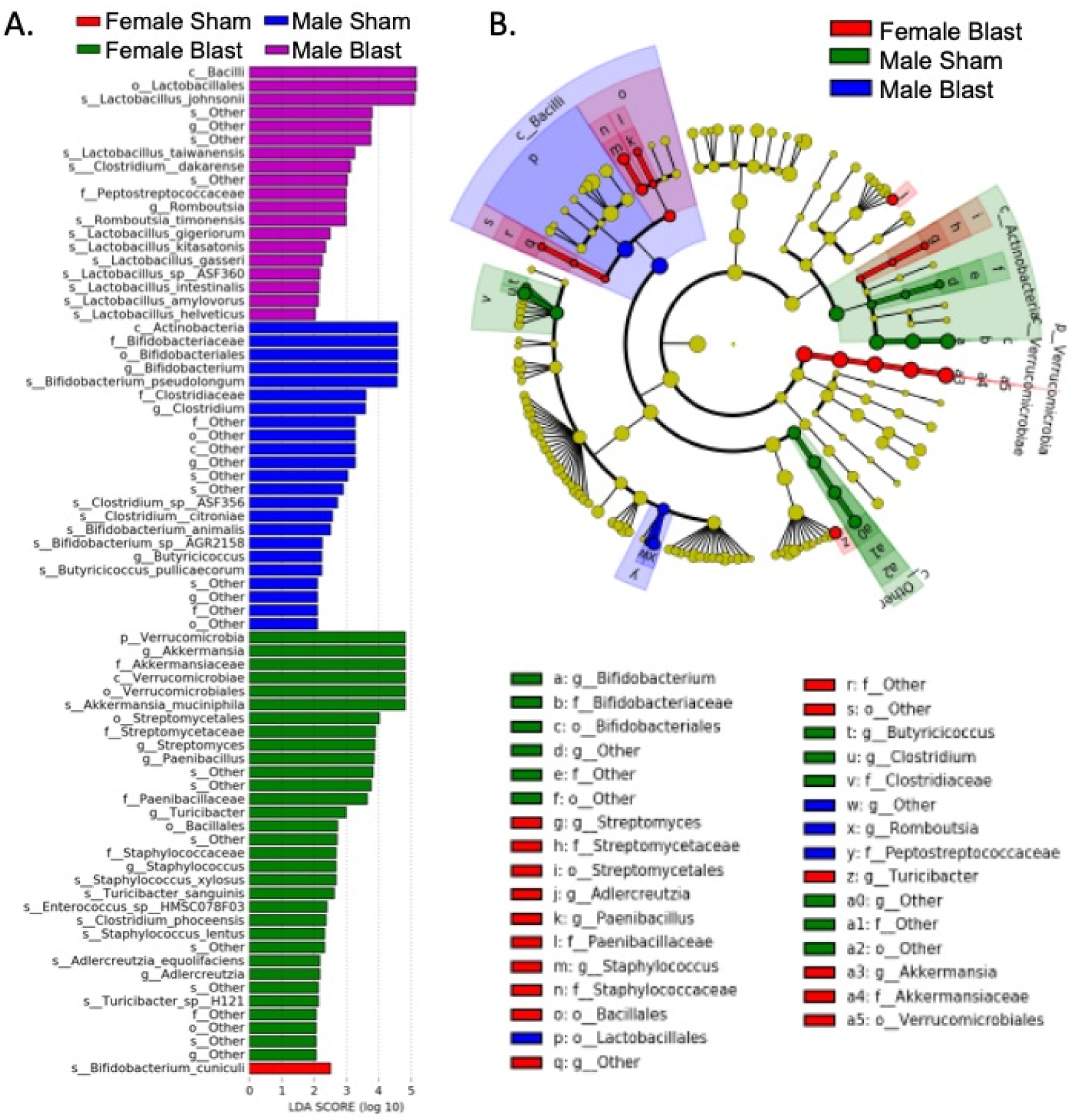
LEfSe analysis of the gut microbial taxonomy. (A). Enriched species (LDA score > 2) in female sham (red), female blast (green), male sham (blue), and male blast (purple). (B) Taxonomic representation of statistically and biologically consistent differences in the four groups. Differences are represented by the color of the most abundant class. Circle diameter is in proportion to that taxon’s abundance.

### Repetitive blast exposure decreases locomotion and increases anxiety-like behavior sub-acutely in both male and female mice

To probe potential locomotor deficits and anxiety-like behavior acutely following blast exposure, we tested male and female mice in a large circular open field 48 hours following their last sham/blast exposure. In distance traveled, mice showed a significant main effect of blast (two-way ANOVA: main effect of exposure *F*(1,85) = 43.15, *p*<0.0001) and no significant interaction effect between exposure and sex (two-way ANOVA: interaction effect *F*(1,85) = 2.573, ns) or main effect of sex (two-way ANOVA: main effect of sex *F*(1,85) = 0.158, ns) (Fig 6a). In entries to the center of the open field, mice showed a significant main effect of blast (two-way ANOVA: main effect of exposure *F*(1,85) = 35.71, *p*=0.0001) and a main effect of sex (two-way ANOVA: main effect of sex *F*(1,85) = 6.591, ns), but no significant interaction effect between exposure and sex (two-way ANOVA: interaction effect *F*(1,85) = 0.641, ns) (Fig 6b). In time spent in the center of the open field, mice showed a significant main effect of blast (two-way ANOVA: main effect of exposure *F*(1,85) = 15.05, *p*=0.0002) and a main effect of sex (two-way ANOVA: main effect of sex *F*(1,85) = 8.736, *p*=0.004), but no significant interaction effect between exposure and sex (two-way ANOVA: interaction effect *F*(1,85) = 0.081, ns) (Fig 6c). Finally, in delay to first center entry, mice showed a significant main effect of blast (two-way ANOVA: main effect of exposure *F*(1,85) = 11.57, *p*=0.001) and no significant interaction effect between exposure and sex (two-way ANOVA: interaction effect *F*(1,85) = 1.99, ns) or main effect of sex (two-way RM ANOVA: main effect of sex *F*(1,85) = 0.045, ns) (Fig 6d).

**Figure 6.**
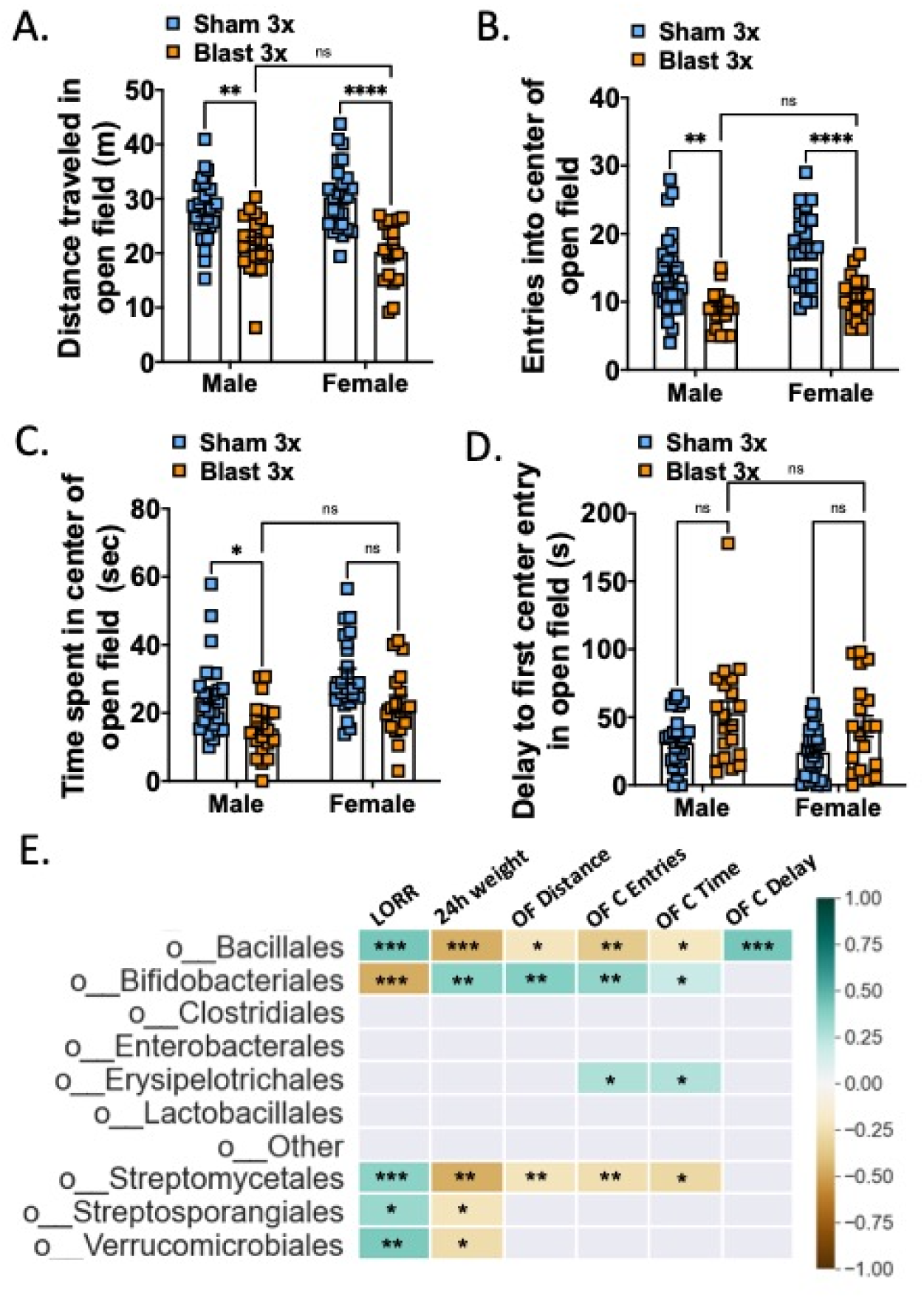
Blast exposure increases acute anxiety-like open field behaviors in both female and male mice. (A) Distance traveled in open field. (B) Entries into center of open field. (C) Time spent in center of open field. (D) Delay to first entry into center of open field. (E) Pearson correlation between bacterial taxa (taxa order that were significantly different between groups from Figure 4) and open field parameters. Two-way ANOVA Šídák’s multiple comparison post-hoc. *p ≤ 0.05, **p ≤ 0.01, ****p ≤ 0.0001, ns = not significant. Values represent mean ± SEM.

To further examine potential interactions between acute behavioral effects and fecal microbiome changes, we computed Pearson correlations and corrected for multiple comparisons using FDR (Figure 6e). The bacterial orders bacillales, bifidobacteriales, and streptomycetales were all correlated with multiple open field parameters as well as LORR and 24-hour weight loss.

### Repetitive blast exposure results in comparable delayed-onset BBB disruption in female and male mice

We previously reported that repetitive (2X or 3X), but not single (1X) blast exposure results in delayed-onset (72 h) BBB disruption to radiolabeled albumin (Logsdon et al., 2018, 2020). Consistent with these prior reports, we found a blast effect on BBB disruption to radiolabeled albumin 72 hours following the last blast exposure (two-way ANOVA: main effect of exposure *F*(1,33) = 5.607, *p*=0.024) but there was no difference between sexes (Figure 7a). Gut microbes and metabolites are capable of modulating BBB function (Logsdon, Erickson, et al., 2018; Parker et al., 2020), thus to further examine potential interactions between BBB disruption and fecal microbiome changes, we computed Pearson correlations and corrected for multiple comparisons using FDR (Figure 7b). Only the bacterial order enterobacterales was correlated with albumin levels after correcting for multiple comparisons.

**Figure 7.**
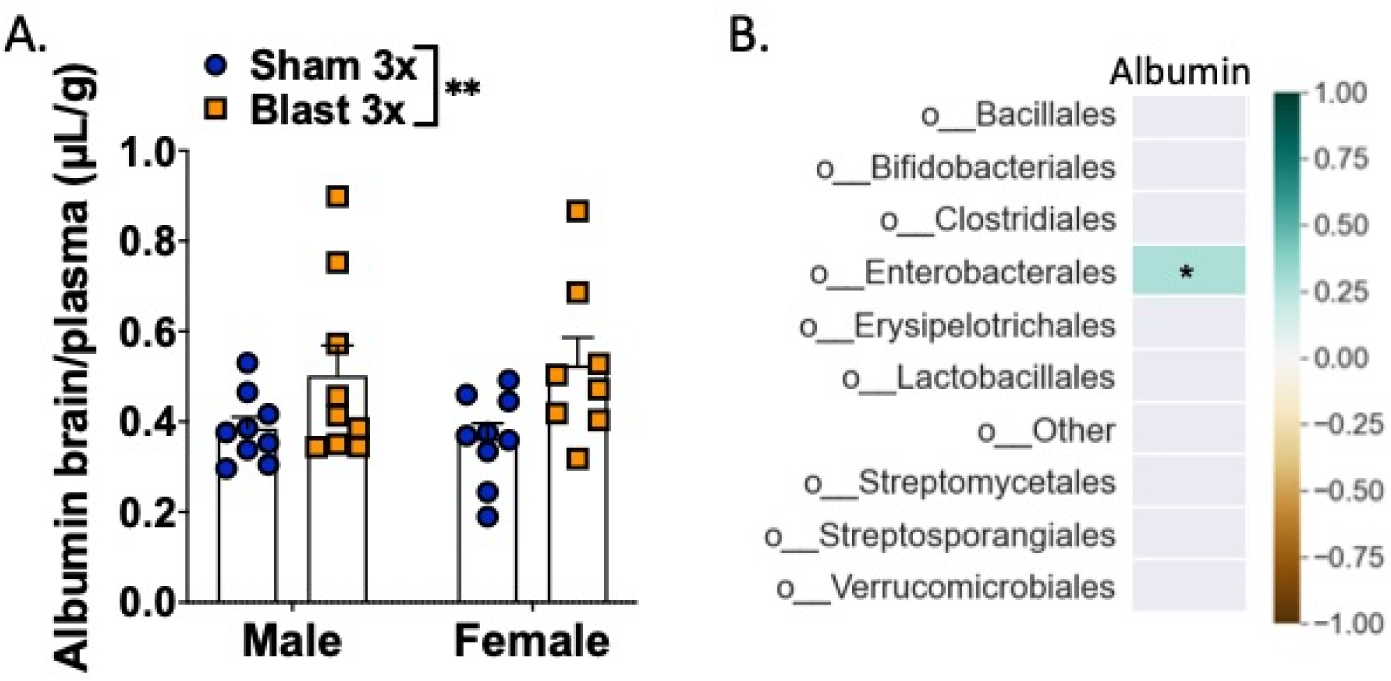
BBB disruption in female and male mice acutely following repetitive blast exposure. (A) Blast increases brain/serum (μl/g) radiolabeled albumin 72 h after final blast. Brain/serum ratios were calculated by dividing the cpm per brain by the cpm per microliter in the corresponding serum and then by the weight of the brain. (B) Pearson correlation between bacterial taxa (taxa order that were significantly different between groups from Figure 4) and albumin BBB permeability. Two-way ANOVA Šídák post-hoc analysis. **p ≤ 0.01, ns = not significant. Values represent means ± SEM and are expressed as microliters per gram of brain tissue.

### Male but not female mice exhibit anxious/aversive-like outcomes one month following repetitive blast exposure

To examine more chronic effects of repeated blast exposure on mTBI and PTSD-like outcomes as have been previously reported in male blast mice (Baskin et al., 2021; Elder et al., 2012; Genovese et al., 2013; Perez-Garcia, De Gasperi, et al., 2018; Perez-Garcia, Gama Sosa, et al., 2018; Schindler et al., 2017; Schindler, Terry, et al., 2021; Statz et al., 2019), one month following their last sham/blast exposure, mice were tested in a behavioral battery consisting of elevated zero maze, acoustic startle, and blast-odorant aversion (Fig 8).

**Figure 8.**
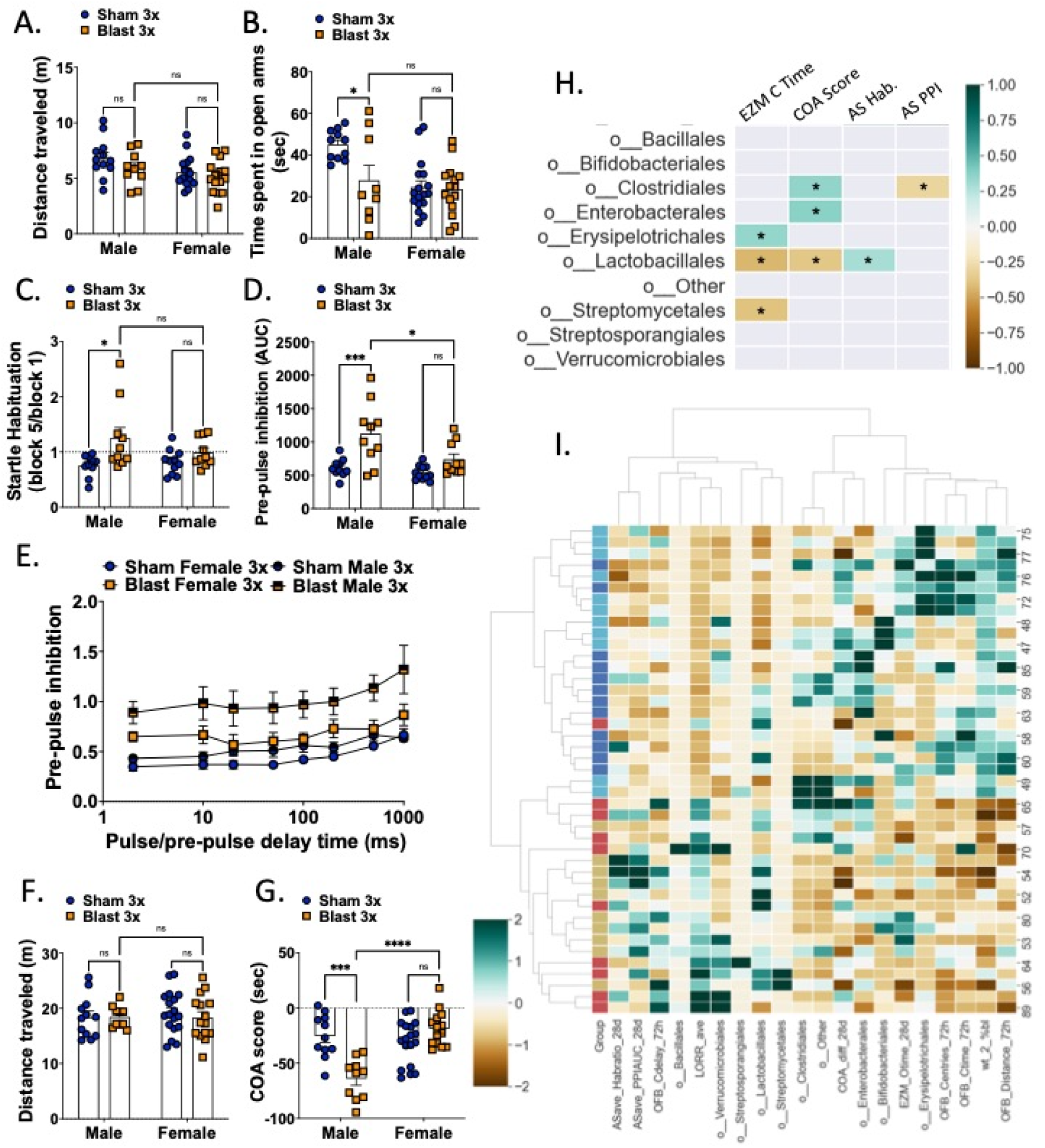
Male but not female mice exhibit chronic blast-induced behavioral deficits. Male and female mice did not differ on total distance traveled on the elevated zero maze 1-month post-blast exposure (A) but male blast mice spent significantly less time than their sham controls in the open arms (B). Only male blast mice showed impaired startle habituation (C) and prepulse inhibition (D, E) on the acoustic startle task. Male and female mice did not differ on total distance traveled in the conditioned odorant aversion posttest (F) but male blast showed an aversion to an odorant previously paired with blast-exposures (G). (H) Pearson correlation between bacterial taxa (taxa order that were significantly different between groups from Figure 4) and chronic behavioral parameters. (I) Heatmap of hierarchical clustering between individual mice vs. behavioral and microbiota composition (taxa order that were significantly different between groups from Figure 4). Each row is a mouse, each column is a parameter. Group column colors: sham female – dark blue; blast female – red; sham male – light blue; blast male yellow. Heatmap colors represent z-score for each parameter computed from all mice. Two-way ANOVA Šídák post-hoc analysis (A-G). *p ≤ 0.05, **p ≤ 0.01, ****p ≤ 0.0001, ns = not significant. Values represent mean ± SEM.

Mice were exposed to an elevated zero maze to probe anxiety-like behaviors with less time spent exploring the open arms indicative of a more anxious phenotype. For distance traveled, there was no significant main effect of exposure (two-way ANOVA *F*(1,48) = 2.518, ns) and no significant interaction effect between exposure and sex (two-way ANOVA: interaction effect *F*(1,48) = 0.209, ns), but there was a significant main effect of sex (two-way ANOVA: main effect of sex *F*(1,48) = 5.95, *p*=0.018) (Fig 8a). Conversely, for time spent in the open arm of the maze, there was a significant main effect of exposure (two-way ANOVA *F*(1,48) = 5.354, *p*=0.025), significant main effect of sex (two-way ANOVA: main effect of sex *F*(1,48) = 10.10, *p*=0.003), and a significant interaction effect between exposure and sex (two-way ANOVA: interaction effect *F*(1,48) = 4.317, *p*=0.043) (Fig 8b). Post-hoc analysis revealed that blast males spent significantly less time in the open arms than sham males (*p*<0.05), whereas female sham and blast mice did not differ in open arm time.

To test for a PTSD-like startle response in the mice, mice went through an acoustic startle session which included assessment of startle habituation and prepulse inhibition (PPI). In the startle habituation test, a decrease in the ability to habituate to a repeated presentation of the same stimulus is representative of a deficit in non-associative learning. In the PPI, a decrease in inhibition by the prepulse is indicative of a difficulty in sensory gating (Valsamis & Schmid, 2011). In line with previous reports in male mice (Baskin et al., 2021), we found a main effect of exposure (two-way ANOVA *F*(1,37) = 8.764, *p*=0.005), but no significant main effect of sex (two-way ANOVA: main effect of sex *F*(1,37) = 0.769, ns) or interaction effect between exposure and sex (two-way ANOVA: interaction effect *F*(1,37) = 2.04, ns) (Fig 8c). PPI was measured following a five minute rest period after the habituation procedure. The area under the curve of the PPI vs. time delay showed a main effect of exposure (two-way ANOVA *F*(1,37) = 17.63, *p*=0.0002) and main effect of sex (two-way ANOVA: main effect of sex *F*(1,37) = 7.074, *p*=0.011), but no interaction effect between exposure and delay (two-way ANOVA: interaction effect *F*(1,37) = 3.218, ns) (Fig 8d). We also examined PPI across time delays and found a main effect of exposure (two-way ANOVA *F*(3,37) = 9.96, *p*<0.0001) and main effect of pulse-prepulse delay time (two-way ANOVA: main effect of delay *F*(5,154) = 13.37, *p*<0.0001) but no interaction effect between exposure and delay (two-way ANOVA: interaction effect *F*(21,259) = 0.688, ns) (Fig 8e). Post-hoc analysis revealed that only blast males demonstrated acoustic startle deficits as compared to sham controls but there was no blast effect in females.

Finally, to test for aversion-like behaviors in mice, prior to every sham/blast exposure, a fresh almond-scented nestlet in a tea diffuser was placed in the home cages of the mice to associate the scent with the experience of the sham or blast trauma. We measured aversion to the almond-scented nestlet in a posttest conducted in a T-maze where the almond scent was placed in the left arm of the T and a clean nestlet with no scent was placed in the right arm of the T. When examining distance traveled, there was no significant main effect of exposure (two-way ANOVA *F*(1,48) = 0.0178, ns), no significant main effect of sex (two-way ANOVA: main effect of sex *F*(1,48) = 0.117, ns), and no significant interaction effect between exposure and sex (two-way ANOVA: interaction effect *F*(1,48) = 0.209, ns) (Fig 8f). Conversely, in the difference between time spent with the odor vs. without the odor (COA), there was a significant main effect of exposure (two-way ANOVA *F*(1,48) = 6.148, *p*=0.017), significant main effect of sex (two-way ANOVA: main effect of sex *F*(1,48) = 14.11, *p*=0.0005), and a significant interaction effect between exposure and sex (two-way ANOVA: interaction effect *F*(1,48) = 24.00, *p*<0.0001) (Fig 8g). Post-hoc analysis revealed that blast males spent significantly less time with the almond odor than sham males (*p*<0.05), whereas females did not differ between sham and blast groups.

To further examine potential interactions between chronic behavioral effects and fecal microbiome changes, we computed Pearson correlations and corrected for multiple comparisons using FDR (Figure 8h). The bacterial orders clostridales and lactobacillales were each correlated with distinct behavioral parameters. Finally, hierarchical clustering across acute and chronic parameters revealed distinct clusters of sham male vs. sham female mice but more mixed clusters consisting of male and female blast mice.

## DISCUSSION

While the effects of repetitive blast mTBI have been documented using rodent models, these studies almost exclusively used male research subjects. Thus, almost nothing is known regarding potential adverse outcomes of repetitive blast exposure in female rodents, resulting in a critical knowledge gap at a time when women are increasingly seen in combat roles with a high probability of repetitive blast exposure. Here we present novel data demonstrating acute, sub-acute, and chronic behavioral and inflammatory changes in male and female mice; intriguingly, while sub-acute behavioral outcomes were relatively consistent between male and female blast mice, chronic adverse behavioral outcomes were only found in male mice. These results highlight the importance of examining potential blast effects across sexes and at multiple time points. Furthermore, differential effects of blast in male vs. female mice in cytokine and gut microbiome changes identify potential diagnostic and therapeutic targets for future development and precision medicine efforts.

Understanding the effects of repetitive blast exposure across sexes is critical for diagnostic and treatment development (Agoston, 2017; Giordano et al., 2020; Gupte et al., 2019; McCabe & Tucker, 2020; Ravula et al., 2022; Simmons et al., 2020; Späni et al., 2018). Increased body armor and improved battlefield medicine make multiple deployments are exceedingly common, greatly increasing the potential for repetitive blast exposure resulting in mTBI comorbid with PTSD (Adamson et al., 2008; Carr et al., 2016; O’Neil et al., 2013; Owens et al., 2008; Simmons et al., 2020; Wenger et al., 2018). Indeed, following blast exposure, mTBI-related persistent post-concussive symptoms (e.g., cognitive difficulties, auditory and vestibular dysfunction, sleep disturbances, negative affect) appreciably overlap with PTSD symptoms (e.g., hypervigilance, aversion to trauma cues, cognitive difficulties, sleep disturbances, negative affect) (Adamson et al., 2008; Elder & Cristian, 2009; Owens et al., 2008; Petrie et al., 2014; Schindler, Baskin, et al., 2021; Schindler et al., 2017). Likewise, both mTBI and PTSD can worsen pre-existing psychiatric disorders such as depression and increase and/or exacerbate substance misuse/addiction and other health risk behaviors (e.g., sensation/novelty seeking, impulsivity, risk taking, irritability/aggression) (María-Ríos & Morrow, 2020; McFarlane, 2010; Miller et al., 2013; Olson-Madden et al., 2012; Schindler, Baskin, et al., 2021; Schindler et al., 2017), potentially compounding negative outcomes following injury and trauma. Indeed, in Veterans we previously reported worse outcomes in relation to ‘risky’ drinking behavior as a function of combat exposure and blast mTBI number (Schindler, Baskin, et al., 2021) and a significant correlation between blast mTBI number and findings on neuroimaging (Meabon et al., 2016; Piantino et al., 2021; Schindler, Baskin, et al., 2021). Results from animal models further highlight the importance of studying repetitive blast exposure (Ahmed et al., 2013; Arun et al., 2020; Heyburn et al., 2019; Kamnaksh et al., 2012; Kawa et al., 2018; Logsdon, Meabon, et al., 2018; Logsdon et al., 2020; Meabon et al., 2016; Schindler et al., 2017; Schindler, Terry, et al., 2021; Schindler, Baskin, et al., 2021); we previously demonstrated PTSD-like outcomes in male mice only following repetitive but not single blast mTBI (Schindler et al., 2021) and more recently demonstrated a blast-dose effect in relation to ethanol sensitivity and ‘binge’-like intake patterns in male mice (Schindler, Baskin, et al., 2021). Likewise, we and others have demonstrated blast-dose effects in relation to biochemical (BBB disruption, cytokine expression) and anxiety-like measures following single vs. repetitive blast exposure (Ahmed et al., 2013; Arun et al., 2020; Heyburn et al., 2019; Kamnaksh et al., 2012; Kawa et al., 2018; Logsdon, Meabon, et al., 2018; Logsdon et al., 2020). Critically, to date there are no reports of repetitive blast TBI research in female rodent models.

Acute increase in pro-inflammatory cytokines and BBB disruption have been well documented in male TBI rodents, but have yet to be investigated in females following repetitive blast trauma. Here we demonstrate robust blast-induced changes in serum and brain cytokine levels that display similar and disparate patterns in male vs. female mice. In line with these results, models of single moderate to severe impact TBI have demonstrated disparate inflammatory outcomes in male vs. female mice acutely following injury (Bromberg et al., 2020; Doran et al., 2019; Krukowski, 2021; Krukowski et al., 2020; Späni et al., 2018; Villapol et al., 2017). Likewise, a recent study using a single mild blast exposure with body shielding reported worse acute and sub-acute neuroinflammatory and BBB outcomes in male vs. female rats (Hubbard et al., 2022). The discrepancy between this study and our current results of similar BBB disruption in male vs. female blast mice is likely due to the repetitive nature of our blast model, suggesting that while female rodents are protected from acute single TBI effects, repetitive blast exposure is sufficient to result in cytokine changes and BBB disruption in females as well as males.

To more fully understand potential differences in adverse outcomes as an effect of sex, we analyzed fecal microbial abundance acutely following repetitive blast exposure. Intestinal (gut) microbiota and the genes they produce (collectively referred to as the gut metagenome) help regulate homeostasis and benefit the host through a range of physiological functions (e.g., protection against pathogens, digestion/nutrient assimilation, regulation of the immune system) (Moloney et al., 2014; Thursby & Juge, 2017). Likewise, the ‘microbiota-gut-brain-axis’ (MGBA) represents a critical bidirectional communication system between the gut and the brain, involving metabolic, endocrine, neural, and immune pathways critical for brain health and cognition (Cryan et al., 2019; Moloney et al., 2014; Thursby & Juge, 2017; Wiley et al., 2017). Conversely, altered microbial abundance and composition (i.e., dysbiosis) can have detrimental effects on health and wellness, including implications for cognitive functioning, mental health, and neurodegeneration (Ceppa et al., 2020; Liu et al., 2020; Ticinesi et al., 2018; Tremlett et al., 2017). Importantly, alterations in gut microbiota have been documented years post injury in individuals with a history of moderate/severe TBI (Brenner et al., 2020; Urban et al., 2020), and preclinical work using animal models also supports the microbiome as playing a mechanistic role in adverse outcomes following stress and trauma (Angoa-Pérez et al., 2020; Matharu et al., 2019; Opeyemi et al., 2021), but no studies thus far have specifically utilized animal models of blast injury (either single or repetitive). In line with our cytokine results, we find similar and disparate effects of repetitive blast on gut microbiome in male vs. female mice. Linear discriminant analysis of fecal samples demonstrated bifidobacteria as significantly increased in sham vs. blast mice, which is already being tested as a potential probiotic for PTSD and depression, raising the possibility of this bacteria as a potential treatment target for further development. Gut microbes and metabolites are also capable of modulating BBB function (Braniste et al., 2014; Logsdon, Erickson, et al., 2018; Parker et al., 2020), and we found the bacterial order enterobacterales correlated with levels of BBB disruption. Enterobacterales belong to the phylum of Proteobacteria, which are potentially involved in both intestinal and extraintestinal disease (Rizzatti et al., 2017).

In relation to potential sex differences in adverse behavioral outcomes following blast exposure, only three studies thus far have been reported (Hubbard et al., 2022; McNamara et al., 2022; Russell, Handa, et al., 2018). Hubbard et al (2022) found increased anxiety-like behavior in male but not female rats in the open field (at 2 days post) and elevated plus maze (at 14 days post) following single blast mTBI with body shielding. Conversely, McNamara et al (2022) found no injury effects in either female or male mice on the elevated plus and zero mazes when tested at 2-4 weeks post single blast exposure. Finally, Russell et al (2018) found increased anxiety-like behavior in male, and to a lesser extent in female, mice at 6 days post single blast exposure. Differences in reported results are likely due to differences in injury severity and specifically the repetitive nature of our injury model. Our behavioral outcomes at the one-month time point are also in line with recent preclinical research looking at animal models of PTSD. One preclinical model of PTSD based on the ability of rats to extinguish fear conditioning, found that female mice were less likely to show long-term fear and anxiety-like behaviors on a variety of behaviors when compared to males with similar deficits in fear extinguishing (Emtyazi et al., 2022).

There are several drawbacks and limitations to our current study, including the lack of brain region specific cytokine measurement and/or biochemical/histochemical assays aimed at understanding potential chances in microglia and astrocytes. Likewise, the behavioral tests conducted were relatively simple and warrant future investigation focused on using more sophisticated operant behavioral paradigms as previously reported in male mice following repetitive blast (Baskin et al., 2021). Finally, we did not measure estrus cycle or gonadal hormone levels, but it is important to note that we consistently find less variability in our female mice as compared to their male counterparts. Likewise, research on sex differences in preclinical models of PTSD repeatedly have indicated no association between estrous stage and development of PTSD-like phenotypes (Emtyazi et al., 2022; Zoladz et al., 2019).

Despite TBI being a leading cause of morbidity and mortality worldwide, as well as a common outcome of modern-day warfare, understanding sex as a biological variable in TBI is still in its infancy (Cogan et al., 2020; Gupte et al., 2019). Results from existing human literature are mixed, and overwhelmingly these reports were not powered to examine potential sex differences. Here we report on a series of translationally relevant outcome measures known to be impacted by TBI in humans, with the goal of providing a survey comparison of male and female mice at acute and chronic timepoint following repetitive blast mTBI. Together, our results demonstrate that adverse effects of repetitive blast are dependent on interactions between gonadal sex, time from injury, biological sample type, and/or behavioral test, and highlight new targets for diagnosis and treatment development aimed at understanding how repetitive blast trauma affects diverse populations.

### Disclaimers

The authors have no conflict of interest to disclose. The views expressed in this scientific presentation are those of the author(s) and do not reflect the official policy or position of the U.S. government or Department of Veteran Affairs.

## Acknowledgements

The authors would like to thank Traci J. Webber, Cindy Pekow, DVM, and Lena Strait-Bodey for their considerable veterinary assistance and care. Timelines were created with BioRender.com. Finally, the authors honor the mice, without which this research would not have been possible.

## Declarations

### Ethics approval and consent to participate

All animal experiments were conducted in accordance with Association for Assessment and Accreditation of Laboratory Animal Care guidelines and were approved by the VA Puget Sound Institutional Animal Care and Use Committee.

### Consent for publication

Not applicable.

### Availability of data and materials

The data in this study are available from the corresponding author upon reasonable request.

## Funding

This work was supported by grants from NIDA Training Grant 2T32DA007278-26 (BMB) a Department of Veteran Affairs (VA) Basic Laboratory Research and Development (BLR&D) Career Development Award 1IK2BX003258 (AGS), a VA BLR&D Merit Review Award 5I01BX002311 (DGC), University of Washington Friends of Alzheimer’s Research (DGC), the UW Royalty Research Fund (DGC), the Jeremy Clark Summer Research Fellowship Award (ESS), and UW NAPE Summer Undergraduate Research Program NIDA DA048736 (KW).

### Authors’ contributions

The work presented here was carried out in collaboration among all authors. AS, BMB, AL, BB, and DG contributed to conception and design of the study. BMB, SJL, AL, BDF, and AS collected and analyzed data. BMB and AS wrote the first draft of the manuscript. All authors contributed to manuscript revision, read, and approved the final manuscript.

